# 3D spatial distribution of *Sost* mRNA and Sclerostin expression in response to *in vivo* mechanical loading

**DOI:** 10.1101/2024.09.23.614612

**Authors:** Quentin A. Meslier, Jacy Hoffmann, Robert Oehrlein, Daniel Kurczy, James R. Monaghan, Sandra J. Shefelbine

## Abstract

Bones adapt to external mechanical loads through a process known as mechanoadaptation. Osteocytes are the bone cells that sense the mechanical environment and initiate a biological response. Investigating the changes in osteocyte molecular expression following mechanical loading has been instrumental in characterizing the regulatory pathways involved in bone adaptation. However, current methods for examining osteocyte molecular expression do not preserve the three-dimensional structure of the bone, which plays a critical role in the mechanical stimuli sensed by the osteocytes and their spatially controlled biological responses.

In this study, we used WISH-BONE to investigate the spatial distribution of *Sost*-mRNA transcripts and its encoded protein, sclerostin, in 3D mouse tibia midshaft following *in vivo* tibia loading. Our findings showed a decrease in the percentage of *Sost*-positive osteocytes predominantly at 25% and 37% of the bone length, and in the posterior-lateral side of the tibia after loading. Sclerostin-positive osteocytes in the loaded legs were found to be similar to the contralateral legs after 2 weeks of loading.

This work is the first to provide a 3D analysis of *Sost* and sclerostin distribution in loaded versus contralateral mouse tibia midshafts. It also highlights the importance of the bone region analyzed and the method utilized when interpreting mechanoadaptation results. WISH-BONE represents a powerful tool for further characterization of mechanosensitive genes regulation in bone and holds potential for advancing the development of new treatments targeting mechanosensitivity-related bone disorders.

## Introduction

Osteocytes are the managers of the adaptation response in bone. They are responsible for sensing the external mechanical stimuli applied to the bone and initiating a biological response, resulting in changes to bone homeostasis [1]. Investigations of osteocytes molecular expression in response to loading have provided greater understanding of the molecular pathways involved in mechanoadaptation [2, 3]. Current methods to analyze gene expression in bone cells require homogenization (RNA-Seq and rt-PCR), or sectioning of the bone (In Situ Hybridization), losing 3D spatial information. Under compression, bones mechanical environment varies along the bone length due. For example, the posterior side of the mouse tibia is in compression whereas the anterior side is in tension during tibial compression [1], [2], [3]. Moreover, the magnitude of mechanical stimuli (strain, fluid flow, and others) spatially varies in the bone which may locally regulate the biological response. Spatial heterogeneity is challenging to capture using tissue sections or methods extracting the molecules of interest out of the samples as they fail to maintain the 3D structure of the bone.

Sclerostin has been shown to play in critical role in bone mechanoadaptation. Sclerostin is released by the osteocytes and prevents bone formation by inhibiting the Wnt/β-catenin pathway. Studies have reported a downregulation of sclerostin expression in the osteocytes 24h after 2 consecutive days of bone mechanical stimulation [1], [4], [5]. An increase in bone formation has been correlated with strain magnitudes and down-regulation of sclerostin expression. In cross-sections, the decrease in osteocytes expressing sclerostin was greater in regions experiencing high strain magnitude, based on finite element simulation [1]. Further studies demonstrated a downregulation of *Sost*, between 3h and 24h after loading, in mouse tibia using RT-PCR [6], RNA sequencing data, and in situ hybridization [7], [8]. In hindlimb unloading models, which reduce the mechanical stimulation in the bone, *Sost* mRNA expression increased, but there was no change in sclerostin protein expression in mouse tibiae [9].

However, mRNA and protein analyses are commonly performed in a whole homogenized portion of the bone or in a few thin 2D sections at specific location of the bone length, limiting the number of cells analyzed. It also reflects the biological response at one specific location, which might not be representative of the biological response in the entire bone and might participate in incomplete interpretation of the results.

The recently developed WISH-BONE methods allows to separately label abundant genes and protein in 3D in adult mouse bones [10]. The WISH-BONE methods enable the investigation of hundreds of thousands of cells in the entire tibia midshaft providing a more complete interpretation of impact of mechanical stimuli on regulation of molecular expression. Until now, investigation of the osteocyte’s response to external loads in 3D was limited due to labeling and imaging challenges.

In this work, we used our recently developed methods, WISH-BONE, to investigate the change in osteocytes expressing *Sost* mRNA in 3D, 24h after uniaxial tibia loading in adult mice. We also investigated the percentage of osteocytes expressing sclerostin after 2 weeks of tibia loading. We hypothesize that regulation of *Sost* expression is dependent on the mechanical environment and therefore spatially varying in 3D with local variations matching known variations in the mechanical environment. We report a decrease of *Sost* mRNA signal in the loaded legs mostly in the posterior-lateral side where the mechanical stimulus is known to be the highest under compression. A decline in protein expression after 2 weeks of loading was captured only at very specific locations. This novel way to explore regulation of mechanosenstive genes and encoded proteins should provide new insights on bone mechanosensitivity and help inform the design of new treatment strategy for bone mass loss.

## Methods

### Animal model

All experiments were approved by Northeastern University’s Institutional Animal Care and Use Committee (IACUC). 22 weeks old female C57BL/6J mice were purchased from Jackson Laboratories. Mice were housed in cages of 5, under normal diet, with a 12-hour dark and light cycle and aged to 23 weeks old. A total of 10 mice were used in this study.

### *In vivo* tibia loading

Mice were placed under anesthesia using 2% isoflurane. The right legs of 23 weeks old mice were loaded using the ElectroForce 5500 (TA instruments, USA). A 10N triangle loading profile was applied over 0.25 seconds, with a load rate of 72 N/s. A 1 second rest period was inserted between each cycle. A total of 100 cycles were applied per loading bout. Left legs were kept as contralateral control for comparison. For the mRNA study, 2 loading bouts were applied on the right legs, 24h and 3h before sacrifice. For the protein study, loading was applied 3 days a week for 2 weeks, which is a commonly used protocol, known to induce bone adaptation measurable with microCT scans. Mice were sacrificed 3 days after the last loading bouts. *In vivo* loading experiment is summarized in Figure 1.

**Figure 1:**
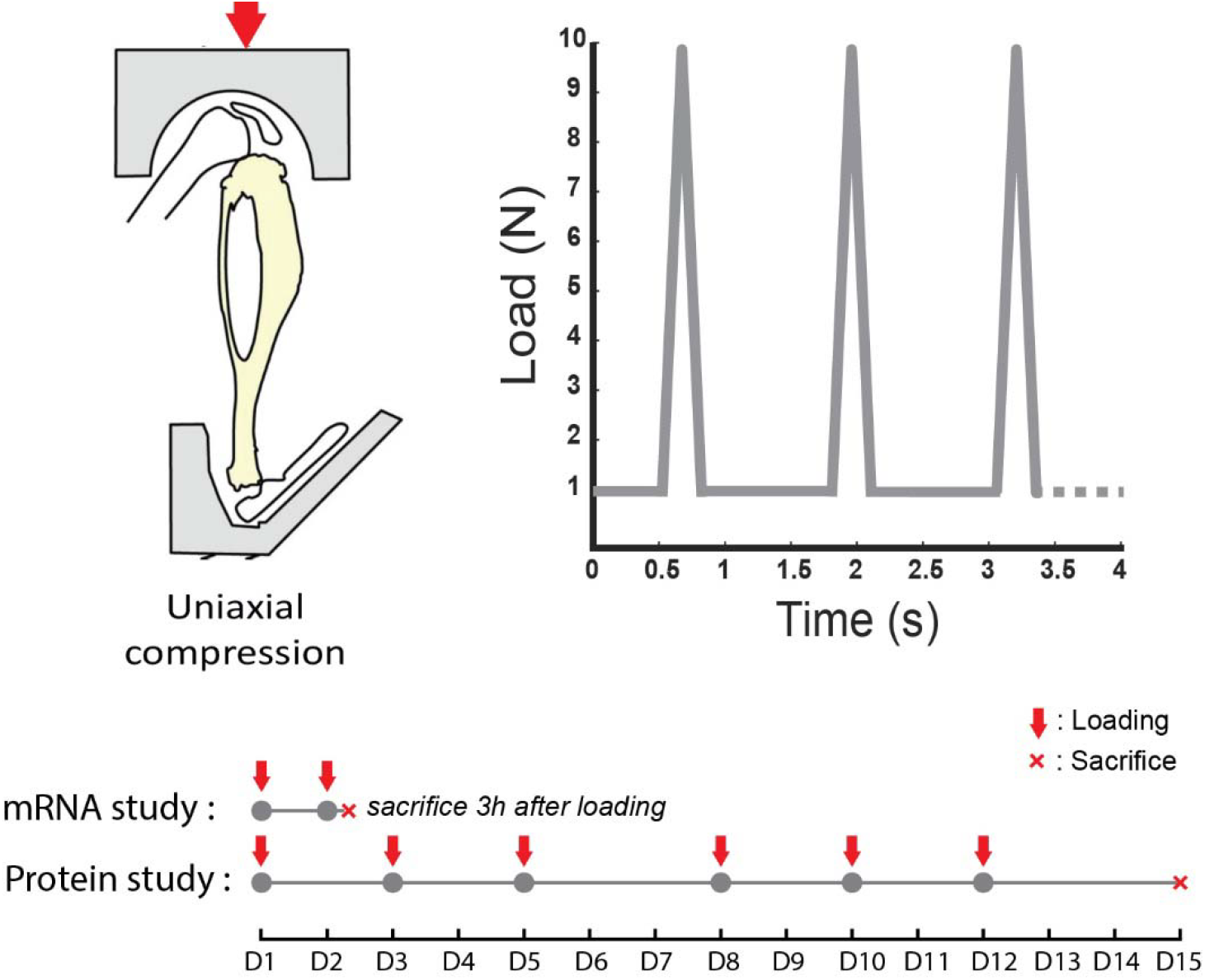
*In vivo* loading experiment summary. The right tibiae were loaded using a uniaxial compression model. 10N triangle loading profile was applied *in vivo* of the mouse tibia. For mRNA analysis, loading was performed 24h and 3h before samples collection. For protein labeling, mice right tibia were loaded 3 days a week for 2 weeks.

### Sample collection, fixation, and decalcification

23 weeks old mice (mRNA study) and 25 weeks old mice (protein study) were sacrificed via CO_2_ inhalation and cervical dislocation. Loaded and contralateral tibiae were collected. External tissues were removed from the bones. The distal end of the tibiae was cut off and the bone marrow was flushed from the proximal end using a 1ml syringe filled with 1xPBS. Samples were then immersed in 4% iced-cold paraformaldehyde (PFA) for 24h at 4⍰c with shaking for fixation. The fixative solution was washed 3 times using 1xPBS and bones were decalcified in 10% EDTA for 2 weeks at 4⍰c with shaking. EDTA was replaced every 3 days. After decalcification, samples were washed in water and store in PBS at 4^⍰^C.

### 3D mRNA labeling

The mRNA labeling protocol was presented in detail in our previous WISH-BONE method and was adapted from previously described protocols [11], [12]. We outline the method briefly here.

Once decalcified, samples were dehydrated using multiple 15min incubation steps in increasing concentration of methanol (MeOH): 25%, 50%, 75%, 100%, and 100% MeOH prepared in 1xPBS with 1% Tween (PBST) and stored in -20^⍰^C overnight. Samples were then rehydrated via successive bath of methanol of decreasing concentration (75%, 50%, 25% methanol in PBST) followed by two 15 min incubations in 100 % PBST at room temperature (RT). Samples were briefly permeabilized in proteinase k (10 µg/ml) for 15 min at RT, post-fixed with 4% PFA for 20 min, and washed 3 times in 1xPBS. Samples were washed in hybridization buffer for 5 min and then prehybridized for 30 min at 37°C. Probe solutions were prepared by diluting the probes at 1:200 in 30% formamide hybridization buffer to a final concentration of 5 µM. Samples were incubated in probe solution overnight at 37°C for hybridization. Preheated 30% formamide wash solution was used to wash out unbound probes by incubating samples 4×15min at 37°C. Samples were then further washed using 5x saline-sodium citrate with 10% Tween (5xSSCT). Samples were incubated in amplification buffer at RT for 30min. DNA hairpins were preheated at 90^⍰^C for 1min, 30sec to ensure proper DNA folding prior to applying to initiators. Hairpins are then cooled to RT for at least 30 min. Hairpins solution was prepared in amplification buffer at a final concertation of 60nM. Samples were incubated in hairpins solution overnight protected from the light. Finally, samples were washed using 5xSSCT and nuclear staining was performed by incubation of the sample with Oxazole Yellow in 1xPBS (1:1000).

To validate labeling, control samples were incubated only with fluorescent DNA hairpins but no DNA probes, which helped quantifying potential signal from non-specific bindings of the fluorescent hairpins in the samples.

### 3D protein labeling

Using the WISH-BONE method [10], decalcified samples were preserved using SHIELD (LifeCanvas Technologies, Cambridge). First, an overnight incubation, at 4°C, in 5ml SHIELD OFF solution (50% SHIELD-epoxy, 25% SHIELD Buffer, and 25% water) was performed to let the epoxy diffuse in the samples. Next samples were incubated in SHIELD ON solution combined with 1% SHIELD-epoxy at 37°C overnight. The SHIELD ON solution allowed the epoxy to polymerize and preserve the sample, while adding 1% of SHIELD-epoxy ensured the good preservation of the bone surfaces. The following day, samples were washed 3 times in 1xPBS to complete samples preservation. Once preserved samples were permeabilized using collagenase. Collagenase solution (10mg/ml) was preheated at 37^⍰^C for 30 min. (Collagenase was aliquoted in 5 ml tubes, stored in the freezer). In the meantime, samples were incubated in preheated 1x HBSS for 10 min at 37°C. Incubation in collagenase solution lasted 6h. To stop the collagenase activities, samples were briefly rinsed in 1xPBS and fixed PFA for 15min. The fixative agent was washed 3 times for 5 min using 1xPBS. Samples were then incubated in PBST with 10% Normal Donkey Serum (NDS) overnight as a blocking step to prevent non-specific binding of the antibodies. Sclerostin was labeled using Goat antibody prepared in PBST solution with 5% NDS and incubated overnight at RT. Samples were then washed 3 times for 10 min in PBST followed by an overnight incubation in PBST. Secondary antibodies (Donkey anti-Goat IgG) were used and prepared in secondary sample buffer with 5% NDS. Samples were incubated for 24h a RT. Oxazole Yellow (1:1000) was added to the secondary antibody solution for nuclear staining. Finally, samples were washed in 1xPBS overnight at RT. To guarantee labeling stability, labeled samples were incubated in 4% PFA for 30 min at RT and washed 3 times in 1xPBS.

Negative control samples were first incubated with Isotype control Goat IgG followed by the same secondary antibodies used in the experimental conditions (Donkey anti-Goat IgG).

### 3D Imaging

Labeled samples were immersed overnight at room temperature in EasyIndex (RI=1.52) for refractive index matching. Next, samples were mounted in 1.5% agarose gel mixed with EasyIndex. Mounted samples were incubated in EasyIndex overnight before imaging. Images were acquired via lightsheet microscopy (SmartSpim, LifeCanvas Technologies, Cambridge) at 3.6x (1.8 µm x 1.8 µm x 2 µm voxel size) and using the following laser wavelength-emission filters combination: 488nm - Chroma ET525/50m, 561 nm - Chroma ET600/50m, and 647 nm - Chroma ET690/50m.

### 3D Cell detection and cell count analysis

TIFF files containing lighthseet images were converted to Zarr files before input to an automated cell detection model (LifeCanvas Technologies, Cambridge, MA). The cell detection pipeline assessed the 3D coordinates of detected cells’ centroid. Performance of the cell detection model was compared to ground truth manual detection and detailed in previous work [10]. F1-score was assessed to be similar to state-of-the-art cell detection model [13]. Images were manually segmented to isolate the cortical bone and remove any remaining external tissues or regions containing bone marrow. To normalize samples length, proximal and distal ends of the bone were segmented out. On the proximal end we used the proximal tibial crest as a landmark, region of the bone located 1mm proximally to the tibial crest was segmented out. Bone located distally to the distal fibula tibia junction was segmented out. As a result, the region of interest analyzed is located between 15% to 55% of the total tibial length. Then a mask of the segmented cortical bone was generated and applied on the cell detection coordinates to exclude cells outside the cortical bone. The total cell counts were based on cell nuclei detection.

To report the number of cells along the bone length, samples were discretized in 20 µm thick sections using MATLAB (Mathworks). The number of cells detected in these sections, based on the nuclear staining, was plotted along the bone length. The number of positive cells for a specific marker was determined by the ratio of cell detection in the marker channel and the total number of cells detected based on the nuclear staining.

To facilitate visualization of the 3D data, the percentage of cells expressing a marker was also reported on two-dimensional surface color maps. To do so, the bone was discretized in quadrants: posterior-lateral, lateral-anterior, antero-medial, and medial-posterior (Figure 3-D). To flatten the 3D point clouds of detected cells, first a centerline was determined by calculating the centroids of 50 cross-sections along the bone. Then cells coordinates were expressed as a radial distance from the centerline and the angle between the radial vector and a reference vector. In the radial direction, the data was average to mimic an orthogonal projection. Coordinates were interpolated on a mesh of 300 points along the Z-axis and 200 points around the angular axis. On the 2D maps, each column corresponds to a quadrant. Samples were rotated along the centerline to guarantee a similar orientation between samples allowing location comparison. The relative change in *Sost*-positive cells between loaded and contralateral legs was reported on those 2D color maps. The relative change was calculated for each mouse by comparing loaded leg and corresponding contralateral leg. The relative change was then averaged into one color map.

In the literature, differences in the number of target-positive cells between experimental conditions are commonly performed in histological sections at specific locations of the bone lengths [1], [4], [9]. For comparison with the literature, we quantified the percentage of *Sost* and sclerostin positive cells present in sections at 25, 37, 45 and 52% of the bone length. We performed the analysis in 20 µm to 400 µm thick simulated bone cross-section to investigate the potential the influence of the selected regions on the quantification results between groups. The mean percentage of target-positive cells of each sample were plotted as individual dots (Figure 4-B).

In addition, we performed a comparison of the percentage of sclerostin-positive cells located specifically in the posterior-lateral region at 37% of the bone length. Three 10 µm-thick sections per sample were analyzed. The number of cells expressing nuclear and sclerostin labeling were manually counted, as per the standard region and method for immunohistochemistry [5], [6], [9].

### Statistical analysis

Statistical significance differences in the number of cell detected between loaded and contralateral groups were first tested using non-parametric 1D Statistical Parametric Mapping (spm1d) [14], two-tailed two-sample t-tests. Statistical Parametric Mapping employs topological interference to identify spatial changes in response to an experimental factor [15]. Spm1d allows a continuous statistical analysis of a 1D data set along the tibia length [14]. Because the results of the normality tests suggested that part of the data was not normally distributed; we used non-parametric spm1d. The null hypothesis assumed that the mean percentage of *Sost*-positive or Sclerostin-positive cells is equivalent between loaded and contralateral groups, for α=0.05. Spm1d test results are presented as SnPM{t} curves along the tibial region of interest for each experimental condition. Red dotted lines indicate the critical thresholds (noted t*) beyond which the null hypothesis is rejected at a Type I rate of α= 0.05.

For comparison with the literature, we statistically tested for significant differences, between loaded and contralateral groups, at 25, 37, 45 and 52% of the bone length using Wilcoxon sum rank test, for α= 0.05. The null hypothesis assumed that the mean percentage of cells expressing the target at the tested location is equal between loaded and contralateral groups. Similarly, the Wilcoxon sum rank test (α= 0.05) was used to test differences in percentage of sclerostin-positive cells in the posterior-lateral region, between loaded and contralateral groups.

## Results

In this work, we investigated the spatial regulation of *Sost* mRNA transcript expression and sclerostin expression in osteocytes following *in vivo* tibia loading. To do so, we used the recently published WISH-BONE method [10] and uniaxial tibia loading [16], [17]. Investigating the spatial change in osteocytes expressing *Sost* mRNA transcript provides indication of the early biological response to compressive loading while expression of sclerostin protein provides a downstream (later in time) response.

### mRNA

We first performed a bulk analysis of the 3D mRNA labeling by considering all the cells detected in the entire sample as one data set. Bulk analysis showed a total number of cells detected based on nuclear staining of about 350,000 cells on average per sample. The total number of cells detected between the loaded and contralateral groups were similar (p=0.6628). However, the total number of cells detected as *Sost*-positive were significantly lower in loaded tibiae compared to contralateral tibiae (p=0.0098) (Figure 2).

**Figure 2:**
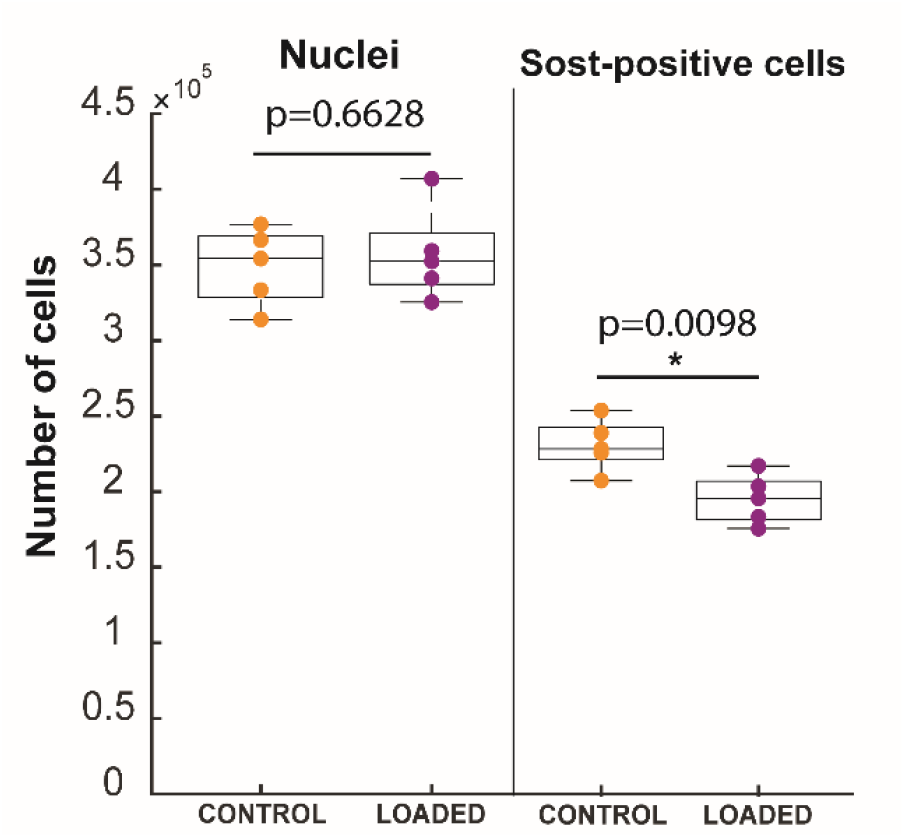
Bulk cell detection analysis of contralateral and loaded samples. Bulk analysis refers to the analysis of cell detection data without considering the spatial information. Wilcoxon sum rank tests were performed to determine if the total number of cells detected were significantly different between loaded and contralateral groups for an alpha value of 0.05.

Figure 3 shows the 3D reconstruction of the lighsheet images for both contralateral and loaded legs labeled for *Sost* mRNA transcripts. Similar to the bulk analysis, the total number of cells was found to be similar along the bone length between both groups (Figure 3-B). The percentage of positive cells were defined by the ratio of the number of cells expressing *Sost* to the number of nuclei in the cortical bone. We observed between 55% and 65% of the cells expressing *Sost mRNA* transcripts along the tibia length in both groups. Comparison with negative control (n=1) showed only 2% of the cells expressing non-specific signal, indicating our labelling is specific for *Sost* (Supplementals Figure-1). Figure 3-C & 3-D show the result of the spatial analysis and the number of *Sost*-positive cells along the bone length. We observed a drop in *Sost*-positive cells in the loaded legs compared to contralateral legs between 20-25% and 35-40% of the tibia length, which are typical regions of adaptation reported in the literature [1], [3], [6], [18]. Spm1d analysis was conducted along the samples length to test for statistical significance between loaded and contralateral groups. Spm1d is a statistical analysis that takes into account the data along the bone length; thus, this test is less sensitive to localized changes between conditions. Spm1d results did not show statistical differences for an α-value of 0.05, suggesting that the changes might be localized. However, when considering bone cross-section of 20 µm, Wilcoxon sum rank tests showed significant decrease of *Sost*-positive cells at 25% and 37% of the bone length between loaded a contralateral leg (α-value= 0.05). In 400 µm thick sections, statistical tests showed significant differences between loaded and contralateral groups at 37% and 52% of the bone length, whereas the region at 25% of the bone length was not significantly different (p=0.056). These results suggest that the region selected for analysis influences the measurement and could alter the interpretation of the results. These results suggest that statistical analysis might lead to different conclusions depending on the region of the bone considered for analysis and the utilized methods.

**Figure 3:**
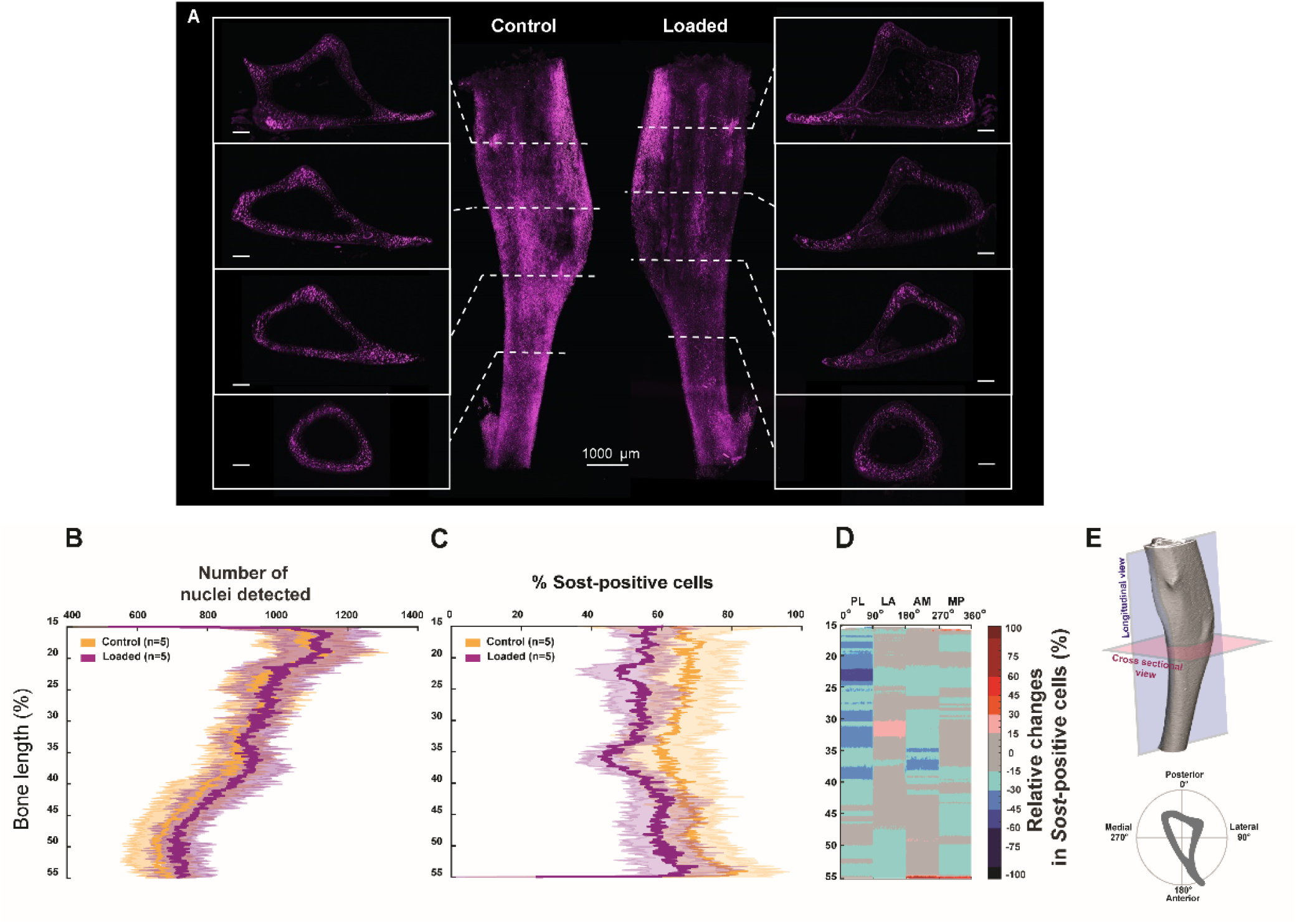
Spatial investigation for *Sost* mRNA expression in loaded vs contralateral legs. A) Lightsheet images of a contralateral and loaded mouse tibia midshafts labeled for *Sost* mRNA using HCR-FISH. The scale bar of the insets represents 200 µm. B) Comparison between loaded and contralateral legs of the total number of cells detected based on nuclear staining along the bone length. Bold lines present the mean percentage of *Sost*-positive cells and shaded bands indicate standard deviation. C) Percentage of *Sost*-positive cells along the bone length of loaded and contralateral legs. D) 2D heat map showing the relative change in *Sost*-positive cells between loaded and contralateral leg along the bone length and within each bone quadrants (PL: Posterior-Lateral, LA: Lateral-Anterior, Anterior-Medial, Medial-Posterior). A negative relative change suggests a decrease in the percentage of *Sost*-positive cells in the loaded leg compared to contralateral leg. E) Illustration showing orientation of the longitudinal plan and cross-sectional plan in the tibia. It also shows the definition of the cross-sectional quadrants used for the 2D heat map.

**Figure 4.**
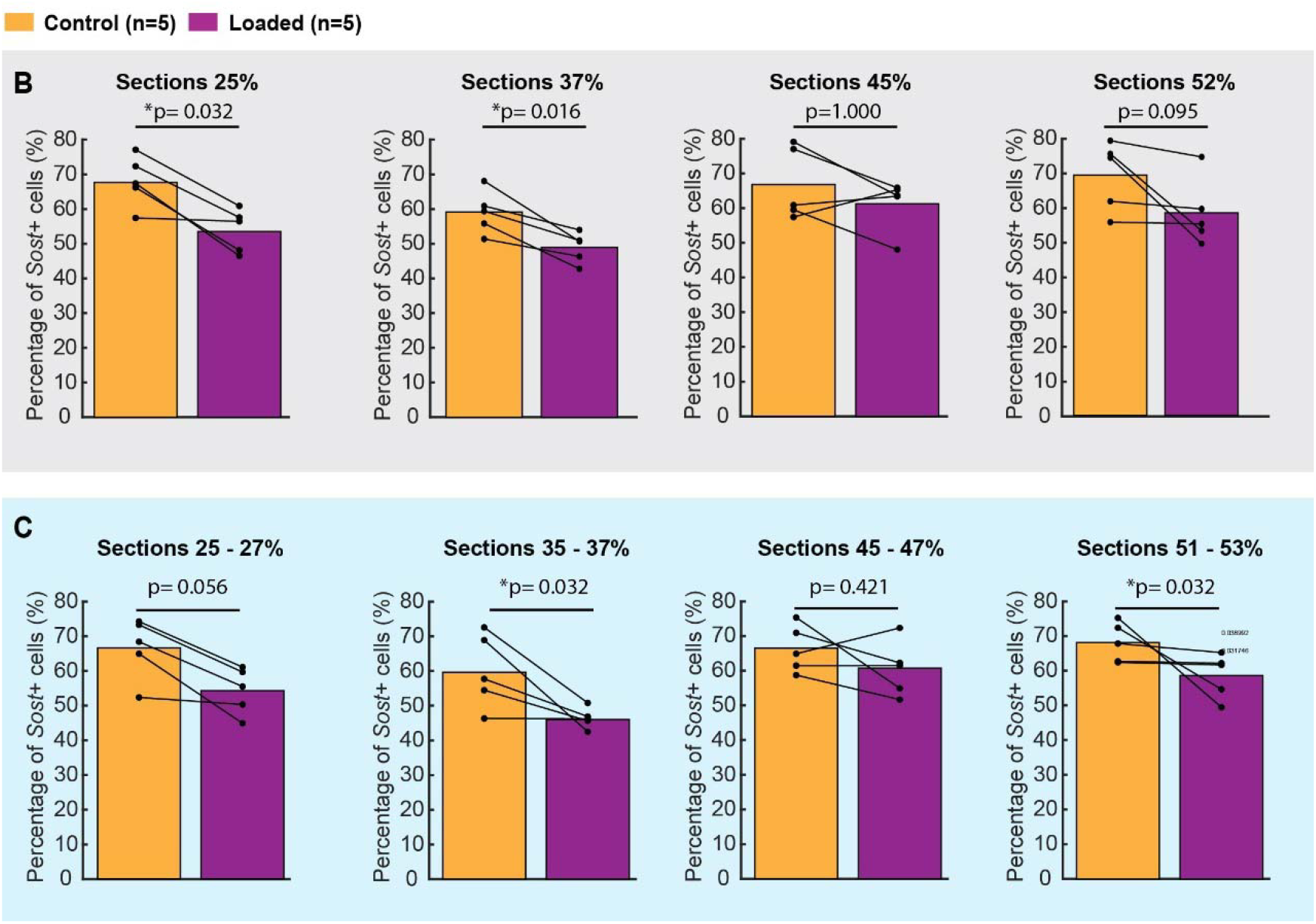
Statistical analysis of *Sost-*positive cells in loaded and contralateral samples. B) Percentage of *Sost*-positive cells detected within 20 µm thick cross-section located at 25%, 37%, 45%, and 52% of the bone length in both groups. Wilcoxon sum rank tests showed a significant difference between loaded and contralateral legs at 25% and 37% of the bone length (α =0.05). Black lines connect loaded and contralateral legs of the same mouse. C) Percentage of *Sost*-positive cells detected within 400 µm thick cross-section located between 25-27%, 35-37%, 45-47%, and 51-53% of the bone length in both groups. Statistical tests showed a significant difference between loaded and contralateral legs in the regions 35-37% and 51-53% (α =0.05).

The 2D colored map showed the relative changes in the percentage of *Sost*-positive cells in the 3D samples (Figure 3-D). A negative relative change suggests a decrease in the number of cells expressing *Sost* in the loaded legs compared to contralateral legs. We observed that most of the decrease in relative change happened in the posterior-lateral side of the bone, where several regions along the bone length presented a decrease in *Sost*-positive cells of 30% or higher compared to contralateral tibiae. The posterior-lateral side of the bone is where the mechanical stimuli are known to be the highest (strain magnitude [1], [19], fluid flow velocity [3], [20], etc.).

### Protein

*Sost* is a gene encoding for the protein sclerostin. As a second step we aimed at investigating the spatial regulation of sclerostin the mouse tibia following loading. Following the same approach, we analyzed sclerostin expression in osteocytes along the bone length after two weeks of loading.

Bulk analysis of the whole-mount protein labeling showed a similar number of total of cells detected in both groups (p=0.7937) based on nuclear staining; as well as a similar number of sclerostin-positive cells in loaded and contralateral groups (p=0.9420). A total of about 380,000 cells on average were detected.

Spatially, we detected a similar number of cells based on nuclei staining along the bone length (Figure 6-B). We observed about 60% of the cells expressing sclerostin along both contralateral and loaded legs. Whereas the percentage of cells presenting non-specific fluorescence, in negative controls, was below 7% (Figure 9). The analysis of the sclerostin-positive cells did not show drops in the number of positive cells along the bone in the loaded compared to contralateral tibia midshafts (Figure 6-C).

The analysis around the bone cross-section, presented as a 2D colored map (Figure 6 – D), did not exhibit a region of change of more than 30%. A few regions had a decrease in the relative change between loaded and contralateral legs around 15%. Those regions were located around 37% and 50% of the bone length, mostly on the posterior-lateral and antero-medial, similar to the mRNA analysis. However, the change in the percentage of sclerostin-positive cells was mild compared to the mRNA analysis.

Spm1d tests showed that loaded and control group were not significantly different in terms of percentage of Sclerostin-positive cells along the bone length (α =0.05). Similar results were found during sections analysis at 25%, 37%, 45%, and 52% of the bone length in 20 µm-thick (Figure 7-B) and 400 µm-thick cross-sections (Figure 7.C).

In the literature, histological analysis of thin 2D sections is used to quantify the number of target-positive osteocytes in the cortical bone between experimental conditions. Here we mimicked this histological approach and quantified the percentage of sclerostin-positive cells in the posterior-lateral region at 37% of the bone length, for both loaded and contralateral samples (Figure 8). Using this method, we quantified an average decrease of 18% in loaded group compared to contralateral (Figure 8.B). The percentage of sclerostin-positive cells was found to be significantly different at this location (α =0.05).

## Discussion

In this study, we used the 3D mRNA and protein labeling method, WISH-BONE, to investigate the spatial regulation of sclerostin expressing and encoding mRNA transcript following *in vivo* tibia loading. Using our spatial analysis, we showed that the method captured a decrease in the percentage of osteocytes expressing *Sost* mRNA transcripts in the loaded group compared to contralateral group. Decreases in the percentage of *Sost*-positive osteocytes were measured around 25% and 37% of the bone length, which are known to be region of adaptation [3], [18], and in the posterior-lateral side of the tibia where the mechanical stimulus is known to be the highest under uniaxial compression [1], [2], [3], [20].

We investigated the percentage of sclerostin-positive cells in the tibia after two weeks of loading. This loading protocol has been shown to induce bone adaptation in mature mouse bone [1], [3], [19], [21]. We found that the percentage of osteocytes expressing sclerostin was similar along the bone length in both experimental conditions. However, significant differences could be found between loaded and contralateral groups when only considering the posterior-lateral region at 37% of the bone length and three 10 um-thick sections per sample.

In this work, we highlight the importance of carefully selecting the region of interest and the influence of the method chosen for its analysis. The 2D color maps (Figure 3.D & 6.D) allow the investigation of molecular regulation around the entire bone cross-section and provide further detail on the spatial location of the regulation compared to simply running the analysis in bulk or along the length of the bone. This method also enables potential correlation between mechanical environments and molecular expression changes [3].

We investigated the regulation of the sclerostin protein in osteocytes in 3D mouse tibia after two weeks of loading. This loading protocol has been commonly used in mouse adaptation studies and is known to lead to bone adaptation in adult mouse bones [1], [3]. However, it was unclear if the expression of sclerostin would be sustained after the two weeks of loading. The percentage of sclerostin-positive osteocytes was similar in loaded and contralateral groups, which differs from previous histological results investigating sclerostin regulation 24h after 2 consecutive days of loading [1], [6], [9]. In our experiment, tibiae were collected 15 days after the start of the loading protocol: 3 days after the last loading session. At this time point sclerostin expression might have returned to baseline in osteocytes. Previous studies showed that the percentage of sclerostin-positive cells in select regions of bone cross-sections returned to baseline 48h after a single loading session [22]. In addition, Holguin et al. found that the number of sclerostin-positive osteocytes was similar in loaded and contralateral groups after 5 days of loading [6]. On the other hand, we know that the repetition of multiple loading cycles and loading sessions is necessary to trigger a bone adaptation response [18], [23] and to obtain measurable amount of bone formation. In addition, 2 weeks of uniaxial tibia loading has been suggested to have long-term/chronic effects on mouse tibia morphology [24] which might have suggested a sustained downregulation of sclerostin in this loading protocol. Because typical histological results quantifying sclerostin-positive cells only focused on thin bone sections at specific location of the bone; one could have hypothesized that sustained downregulation of sclerostin might have been missed at other locations of the bone. Here we reported no changes in percentage of sclerostin-positive osteocytes in the entire 3D tibia midshaft, after two weeks of loading (6 loading sessions of 100 cycles), which suggests the lack of sustained effect of the loading protocol on the downregulation of sclerostin expression in osteocytes.

The potential limitation of this approach is that our labeling and cell detection model might prevent capturing slight downregulation of sclerostin expression in individual cells. A small change in sclerostin expression may result in a minor change in fluorescent due to amplification of the signal through secondary antibodies. These slightly less bright cells might still be bright enough to be detected in our current detection pipeline, preventing assessment of the level of expression. Less-sensitive cell detection models could be trained to investigate this question.

Despite the similarities of the 3D data set between loaded and contralateral groups (Figure 5,6,7), we found statistically significant differences in the posterior-lateral region at 37% of the bone between our groups (Figure 8). At this location, three 10 µm-tick sections per sample were analyzed, as commonly performed in the literature. We measured a decrease of about 18% in sclerostin-positive cells. These results highlight the influences of the method of analysis on the interpretation of the effect of the mechanical loading used. Moreover, it suggests a potential lack of representativity of the 2D histological sections to characterize the mechanoadaptation response in the entire bone.

**Figure 5:**
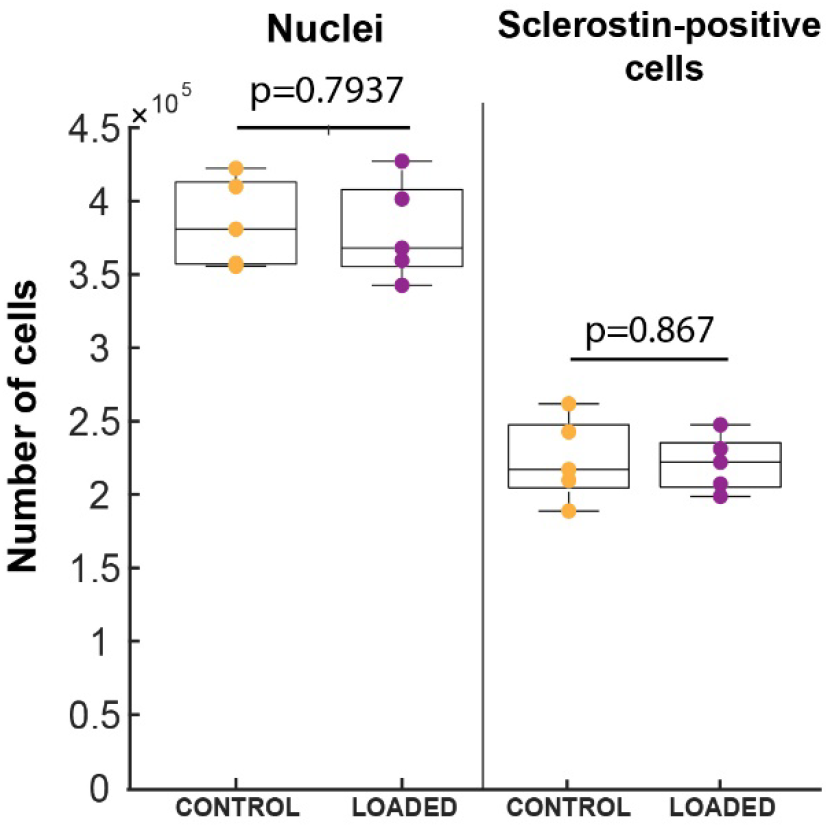
Bulk cell detection analysis of contralateral and loaded samples. Bulk analysis refers to the analysis of cell detection data without considering the spatial information. Wilcoxon sum rank tests (α= 0.05) were performed to determine if the number of cells detected was significantly different between loaded vs contralateral groups.

**Figure 6:**
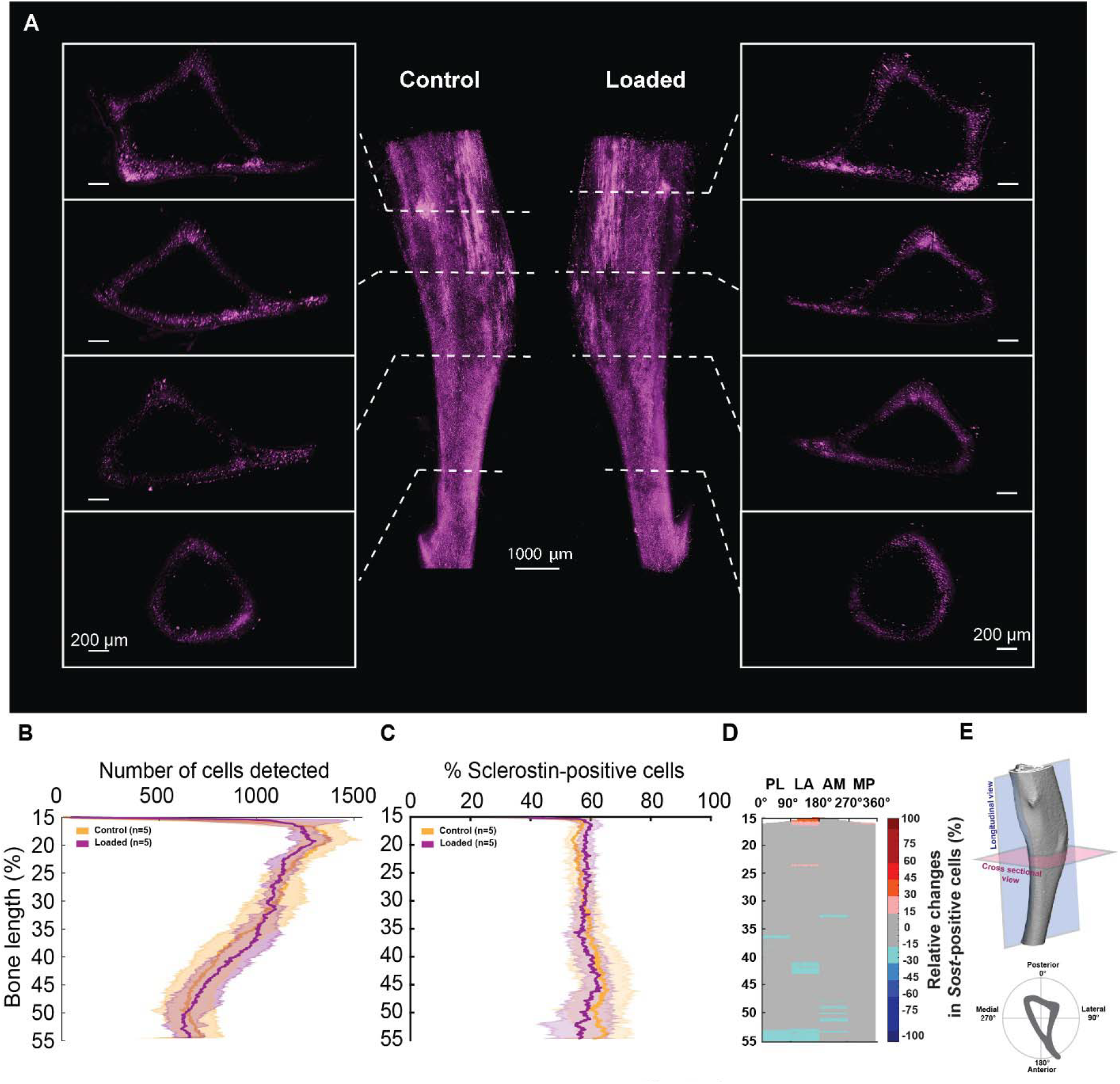
Spatial investigation for Sclerostin expression in loaded vs contralateral legs. A) Lightsheet images of a contralateral and loaded mouse tibia midshafts labeled for Sclerostin using whole-mount immunolabeling. B) Comparison between loaded and contralateral legs of the total number of cells detected based on nuclear staining along the bone length. C) Percentage of Sclerostin-positive cells along the bone length of loaded and contralateral legs. D) 2D heat map showing the relative change in Sclerostin-positive cells between loaded and contralateral leg along the bone length and within each bone quadrants (PL: Posterior-Lateral, LA: Lateral-Anterior, Anterior-Medial, Medial-Posterior). A negative relative change suggests a decrease in the percentage of Sclerostin -positive cells in the loaded compared to contralateral legs. E) Illustration showing the plan the orientation of the longitudinal plan and cross-sectional plan in the tibia. It also shows the definition of the cross-sectional quadrants used for the 2D heat map.

**Figure 7:**
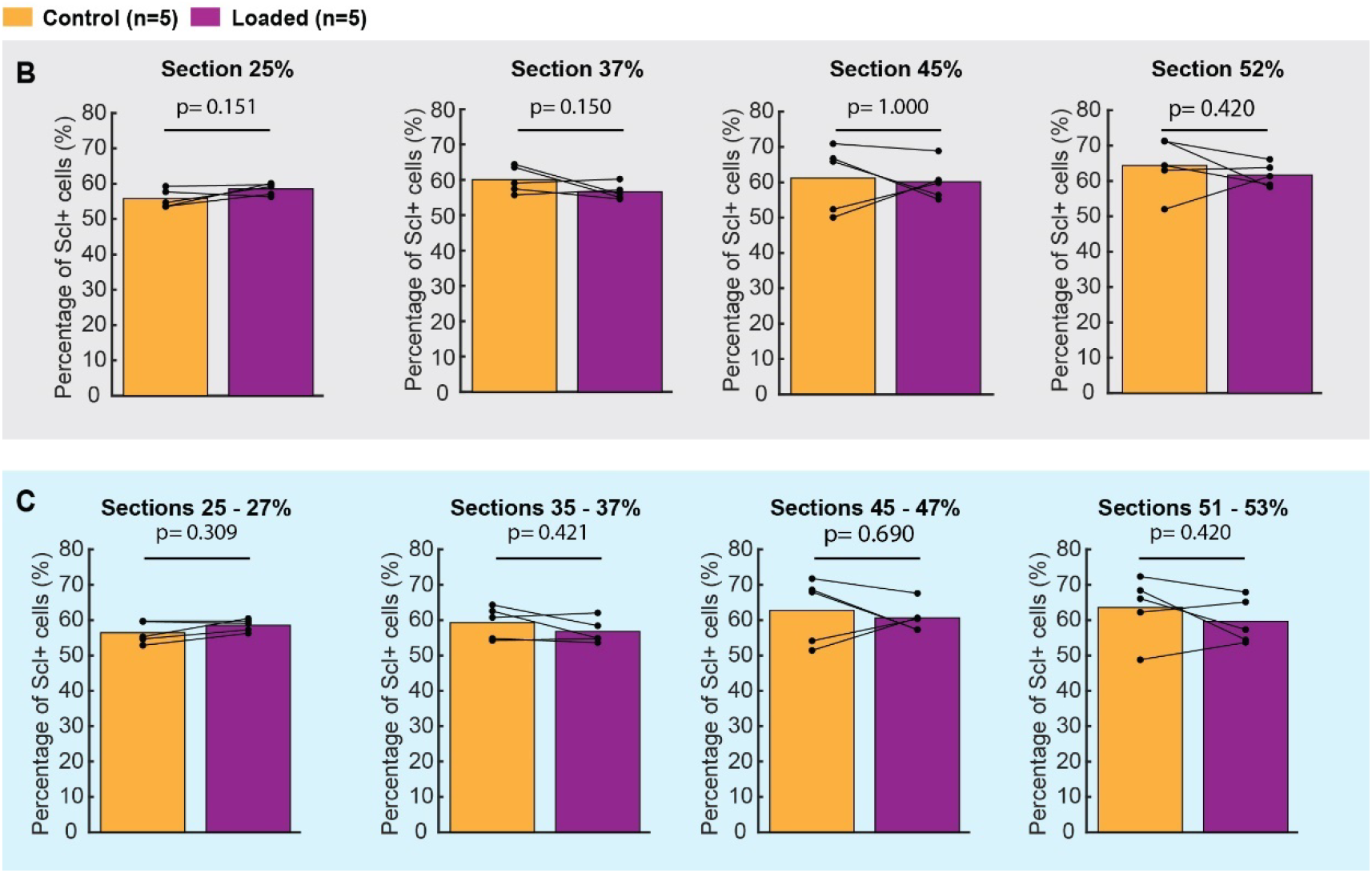
Statistical analysis of Sclerostin-positive *cells*. B) Percentage of Sclerostin-positive cells detected within 20 µm thick cross-section located at 25%, 37%, 45%, and 52% of the bone length in both groups. Solid lines indicate samples from the same mouse. Statistical analysis did not show a significant difference between loaded and contralateral legs at the tested location (α =0.05). C) Percentage of Sclerostin-positive cells detected within 400 µm thick cross-section located between 25-27%, 35-37%, 45-47%, and 51-53% of the bone length in both groups. No significant difference between loaded and contralateral legs were detected (α =0.05).

**Figure 8:**
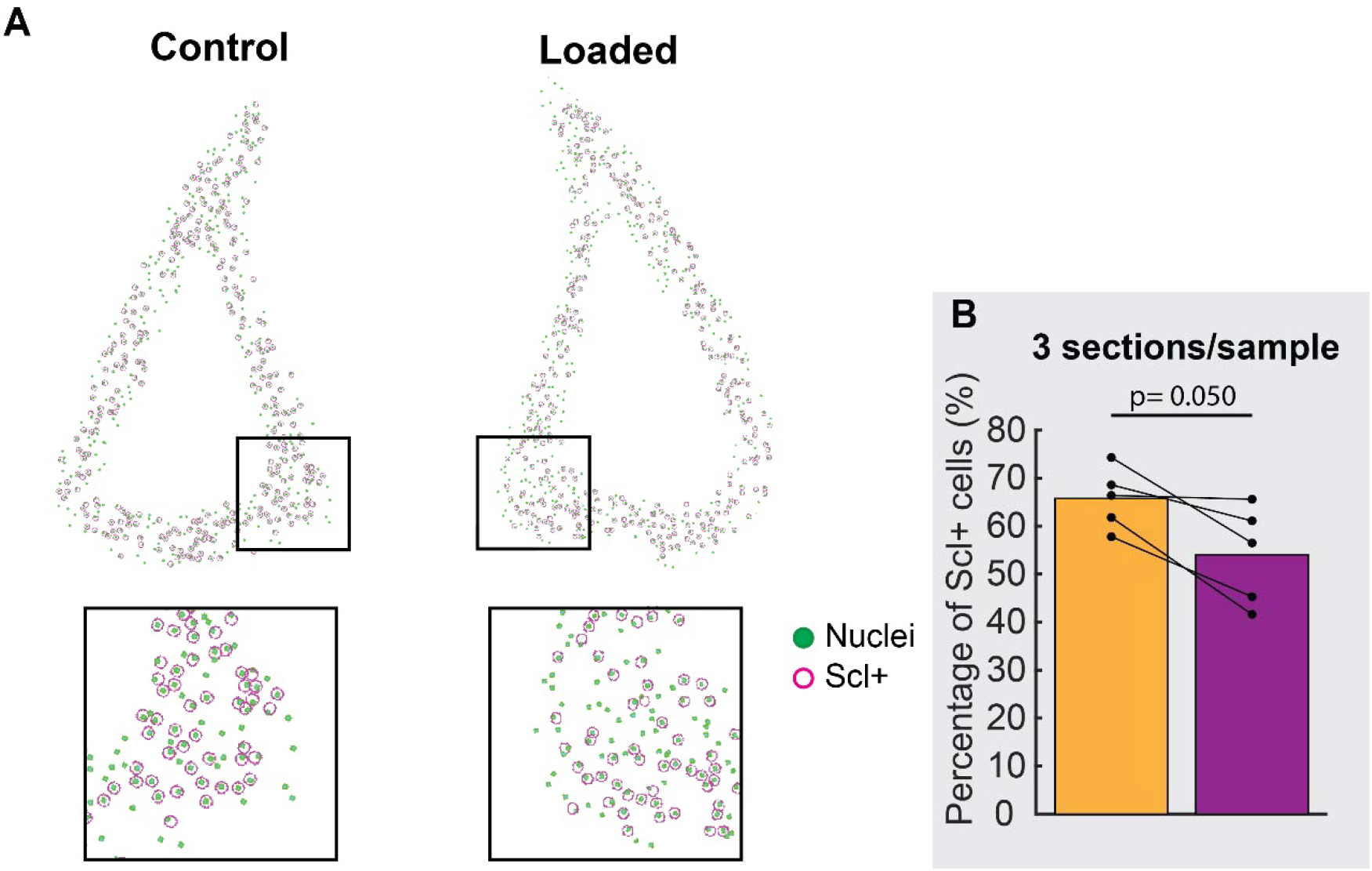
Histological analysis of the loaded and contralateral bones. A) Simulated tibia cross-sections from loaded and contralateral legs based on 3D lightsheet data. Analyzed sections were localized at 37% of the bone length. Cells located in the posterior-lateral regions (insets) of the sections were manually counted and the percentage of cells expressing sclerostin (Scl+) was reported. B) Results were reported after analysis of three consecutive 10 µm-thick sections per sample. Analysis was conducted using the 3D cells detection data from the lightsheet images.

## Conclusions

In this work we investigated the 3D distribution of *Sost* mRNA and sclerostin expression in osteocytes respectively after 24h and 2 weeks of uniaxial tibia compression. The new WISH-Bone method provides important information of the location of the downregulation of *Sost* in 3D. The protein analysis emphasized the influence of the methods and locations of quantification of sclerostin-positive cells on the interpretation of the mechanoadaptation response. Investigating abundantly expressed genes and proteins in 3D should provide a better understanding of the mechanoadpation response in healthy and pathological bones.

## Supporting information

Supplemental Results

## Acknowledgements

This work was funded by the National Science Foundation CMMI # 2010010 and the National Science Foundation INTERN supplement.

We thank the Institute for Chemical Imaging of Living Systems (RRID:SCR_022681) at Northeastern University for consultation and instrument support.

In addition, we thank LifeCanvas technology for their guidance in tissue labeling and imaging.

## Author Contributions

**Quentin A Meslier:** Conceptualization, Methodology, Investigation, Formal analysis, Visualization, Writing – original draft, Writing – review & editing. **Jacy Hoffmann:** Formal analysis. **Robert Oehrlein:** Formal analysis. **Daniel Kurczy:** Formal analysis. **James R. Monaghan and Sandra J. Shefelbine:** Conceptualization, methodology, Writing – review & editing, Supervision, Funding acquisition.

## Reference

[1] A. Moustafa et al., “Mechanical loading-related changes in osteocyte sclerostin expression in mice are more closely associated with the subsequent osteogenic response than the peak strains engendered,” Osteoporos Int, vol. 23, no. 4, pp. 1225–1234, Apr. 2012, doi: 10.1007/s00198-011-1656-4.

[2] H. Razi et al., “Skeletal maturity leads to a reduction in the strain magnitudes induced within the bone: A murine tibia study,” Acta Biomaterialia, vol. 13, pp. 301–310, Feb. 2015, doi: 10.1016/j.actbio.2014.11.021.

[3] Q. A. Meslier, N. DiMauro, P. Somanchi, S. Nano, and S. J. Shefelbine, “Manipulating load-induced fluid flow in vivo to promote bone adaptation,” Bone, vol. 165, p. 116547, Dec. 2022, doi: 10.1016/j.bone.2022.116547.

[4] L. B. Meakin, G. L. Galea, T. Sugiyama, L. E. Lanyon, and J. S. Price, “Age-Related Impairment of Bones’ Adaptive Response to Loading in Mice Is Associated With Sex-Related Deficiencies in Osteoblasts but No Change in Osteocytes: AGE-RELATED IMPAIRMENT OF BONES’ ADAPTIVE RESPONSE TO MECHANICAL LOADING,” J Bone Miner Res, vol. 29, no. 8, Art. no. 8, Aug. 2014, doi: 10.1002/jbmr.2222.

[5] A. Moustafa et al., “The mouse fibula as a suitable bone for the study of functional adaptation to mechanical loading,” Bone, vol. 44, no. 5, pp. 930–935, May 2009, doi: 10.1016/j.bone.2008.12.026.

[6] N. Holguin, M. D. Brodt, and M. J. Silva, “Activation of Wnt Signaling by Mechanical Loading Is Impaired in the Bone of Old Mice: LOAD-INDUCED ACTIVATION OF WNT SIGNALING IMPAIRED IN BONE OF OLD MICE,” J Bone Miner Res, vol. 31, no. 12, pp. 2215–2226, Dec. 2016, doi: 10.1002/jbmr.2900.

[7] C. Chlebek, J. A. Moore, F. P. Ross, and M. C. H. van der Meulen, “Molecular Identification of Spatially Distinct Anabolic Responses to Mechanical Loading in Murine Cortical Bone,” J of Bone & Mineral Res, p. jbmr.4686, Sep. 2022, doi: 10.1002/jbmr.4686.

[8] N. H. Kelly, J. C. Schimenti, F. P. Ross, and M. C. H. Van Der Meulen, “Transcriptional profiling of cortical versus cancellous bone from mechanically-loaded murine tibiae reveals differential gene expression,” Bone, vol. 86, pp. 22–29, May 2016, doi: 10.1016/j.bone.2016.02.007.

[9] A. G. Robling et al., “Mechanical Stimulation of Bone in vivo Reduces Osteocyte Expression of Sost/Sclerostin,” Journal of Biological Chemistry, vol. 283, no. 9, Art. no. 9, Feb. 2008, doi: 10.1074/jbc.M705092200.

[10] Q. A. Meslier, T. J. Duerr, B. Nguyen, J. R. Monaghan, and S. J. Shefelbine, “WISH-BONE: Whole-mount In Situ Histology, to label osteocyte mRNA and protein in 3D adult mouse bones,” Mar. 15, 2024. doi: 10.1101/2024.03.14.585053.

[11] A. M. Lovely, T. J. Duerr, D. F. Stein, E. T. Mun, and J. R. Monaghan, “Hybridization Chain Reaction Fluorescence In Situ Hybridization (HCR-FISH) in Ambystoma mexicanum Tissue,” in Salamanders, vol. 2562, A. W. Seifert and J. D. Currie, Eds., in Methods in Molecular Biology, vol. 2562., New York, NY: Springer US, 2023, pp. 109–122. doi: 10.1007/978-1-0716-2659-7_6.

[12] H. M. T. Choi et al., “Third-generation in situ hybridization chain reaction: multiplexed, quantitative, sensitive, versatile, robust,” Development, vol. 145, no. 12, p. dev165753, Jun. 2018, doi: 10.1242/dev.165753.

[13] D. Kaltenecker et al., “Virtual reality-empowered deep-learning analysis of brain cells,” Nat Methods, vol. 21, no. 7, pp. 1306–1315, Jul. 2024, doi: 10.1038/s41592-024-02245-2.

[14] T. C. Pataky, “One-dimensional statistical parametric mapping in Python,” Computer Methods in Biomechanics and Biomedical Engineering, vol. 15, no. 3, pp. 295–301, Mar. 2012, doi: 10.1080/10255842.2010.527837.

[15] K. J. Friston, Ed., Statistical parametric mapping: the analysis of funtional brain images, 1st ed. Amsterdam1; Boston: Elsevier/Academic Press, 2007.

[16] R. L. De Souza, M. Matsuura, F. Eckstein, S. C. F. Rawlinson, L. E. Lanyon, and A. A. Pitsillides, “Non-invasive axial loading of mouse tibiae increases cortical bone formation and modifies trabecular organization: A new model to study cortical and cancellous compartments in a single loaded element,” Bone, vol. 37, no. 6, Art. no. 6, Dec. 2005, doi: 10.1016/j.bone.2005.07.022.

[17] J. Fritton, E. Myers, T. Wright, and M. Vandermeulen, “Loading induces site-specific increases in mineral content assessed by microcomputed tomography of the mouse tibia,” Bone, vol. 36, no. 6, Art. no. 6, Jun. 2005, doi: 10.1016/j.bone.2005.02.013.

[18] H. Yang, R. E. Embry, and R. P. Main, “Effects of Loading Duration and Short Rest Insertion on Cancellous and Cortical Bone Adaptation in the Mouse Tibia,” PLoS ONE, vol. 12, no. 1, Art. no. 1, Jan. 2017, doi: 10.1371/journal.pone.0169519.

[19] J. Piet, D. Hu, Q. Meslier, R. Baron, and S. J. Shefelbine, “Increased Cellular Presence After Sciatic Neurectomy Improves the Bone Mechano-adaptive Response in Aged Mice,” Calcif Tissue Int, vol. 105, no. 3, pp. 316–330, Sep. 2019, doi: 10.1007/s00223-019-00572-7.

[20] A. F. Pereira, B. Javaheri, A. A. Pitsillides, and S. J. Shefelbine, “Predicting cortical bone adaptation to axial loading in the mouse tibia,” J. R. Soc. Interface., vol. 12, no. 110, p. 20150590, Sep. 2015, doi: 10.1098/rsif.2015.0590.

[21] K. Eller et al., “Mechanoadaptation of the bones of mice with high fat diet induced obesity in response to cyclical loading,” Journal of Biomechanics, vol. 124, p. 110569, Jul. 2021, doi: 10.1016/j.jbiomech.2021.110569.

[22] N. Lara-Castillo et al., “In vivo mechanical loading rapidly activates β-catenin signaling in osteocytes through a prostaglandin mediated mechanism,” Bone, vol. 76, pp. 58–66, Jul. 2015, doi: 10.1016/j.bone.2015.03.019.

[23] D. Sun, M. D. Brodt, H. M. Zannit, N. Holguin, and M. J. Silva, “Evaluation of loading parameters for murine axial tibial loading: Stimulating cortical bone formation while reducing loading duration: EVALUATION OF LOADING PARAMETERS FOR MURINE AXIAL TIBIAL,” J. Orthop. Res., Oct. 2017, doi: 10.1002/jor.23727.

[24] B. Javaheri et al., “Lasting organ-level bone mechanoadaptation is unrelated to local strain,” Sci. Adv., vol. 6, no. 10, p. eaax8301, Mar. 2020, doi: 10.1126/sciadv.aax8301.

