## Supplemental Results for "3D spatial distribution of *Sost* mRNA and Sclerostin expression in response to *in vivo* mechanical loading"

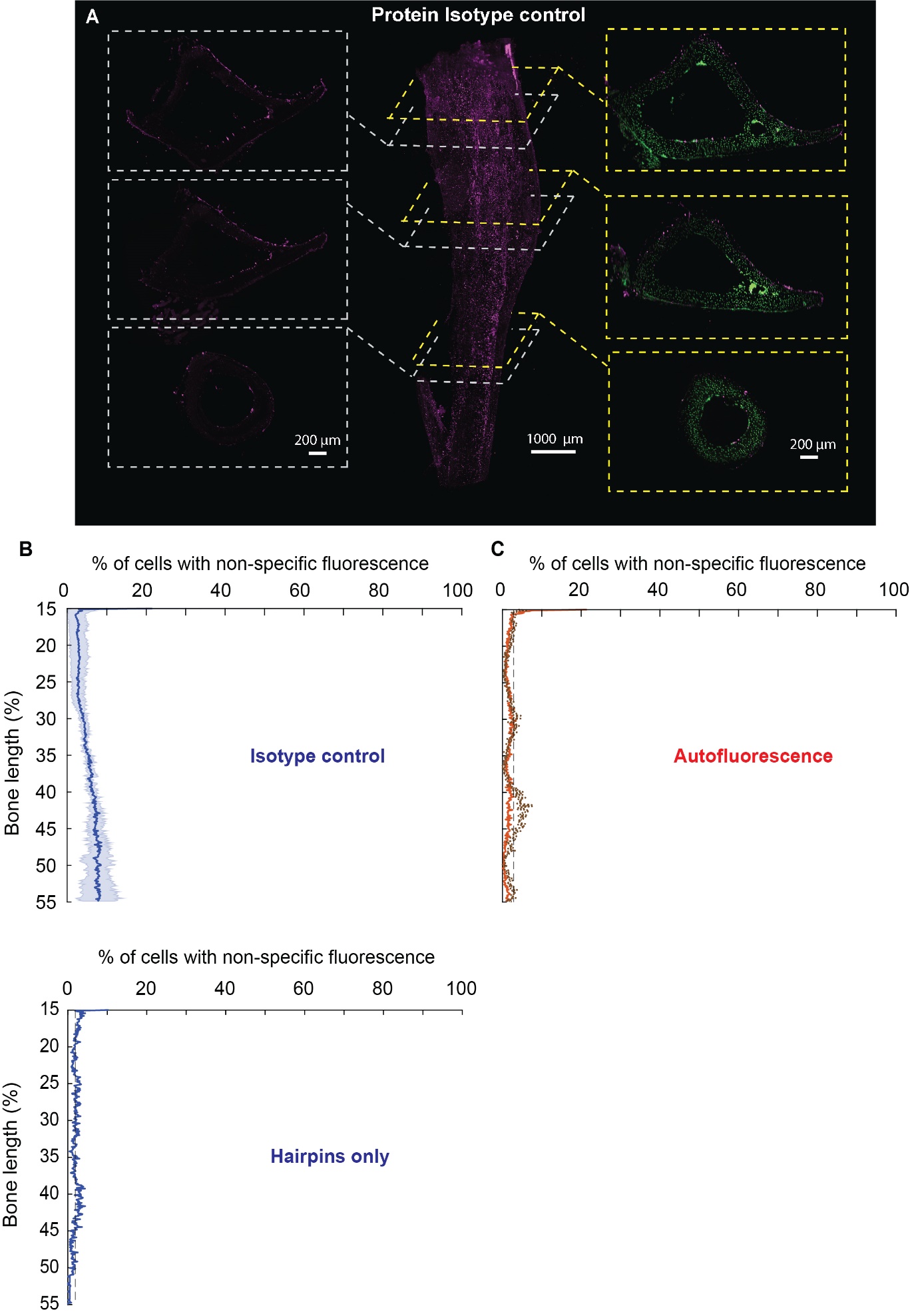

**Supplementals Figure 1:** Spatial investigation for mRNA labeling negative control. The graph shows the percentage cells detected as expressing non-specific fluorescence along the bone length (n=1). Non-specific fluorescence us the results of the incubation with fluorescent DNA hairpins without prior incubation with gene specific probes.

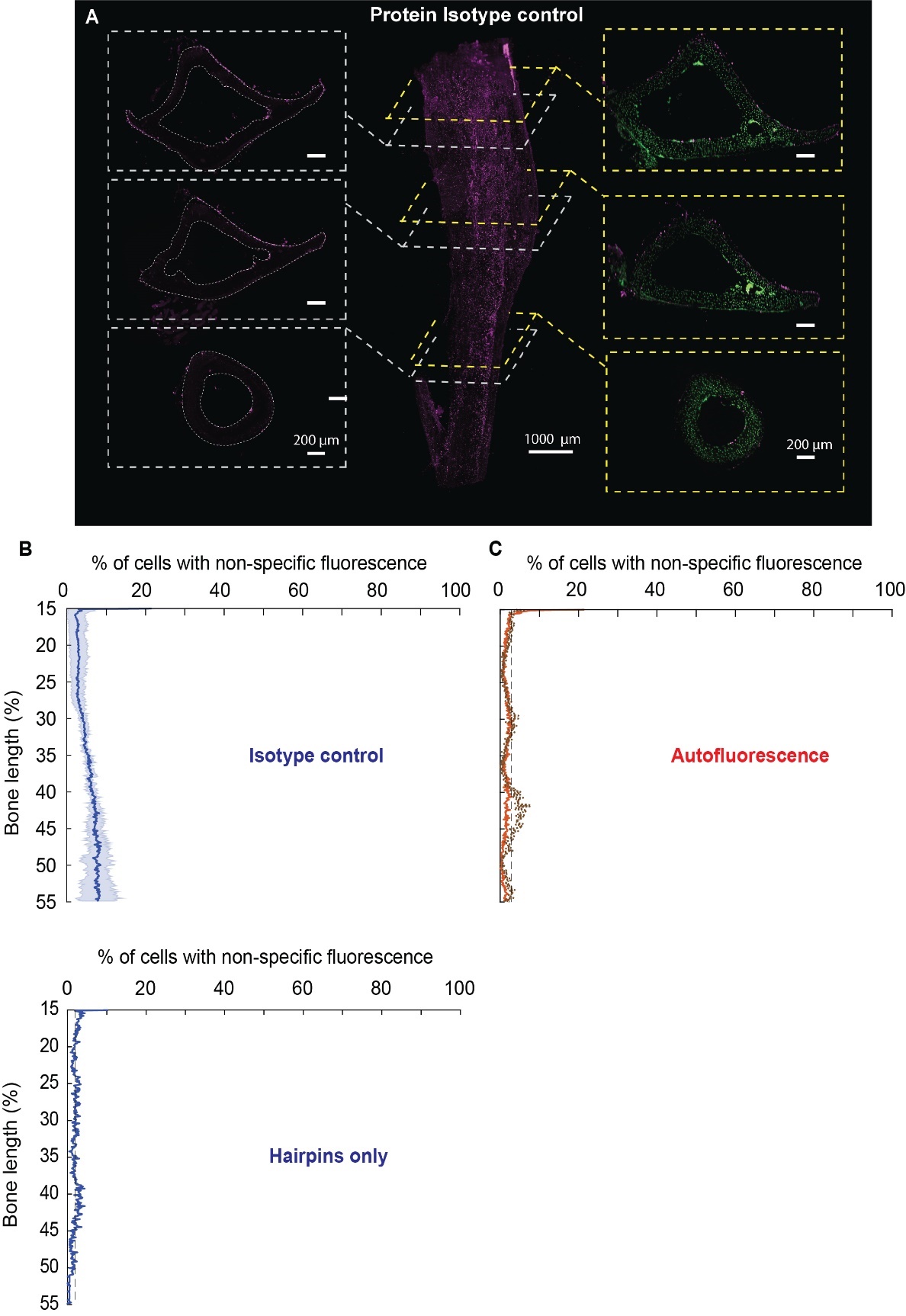

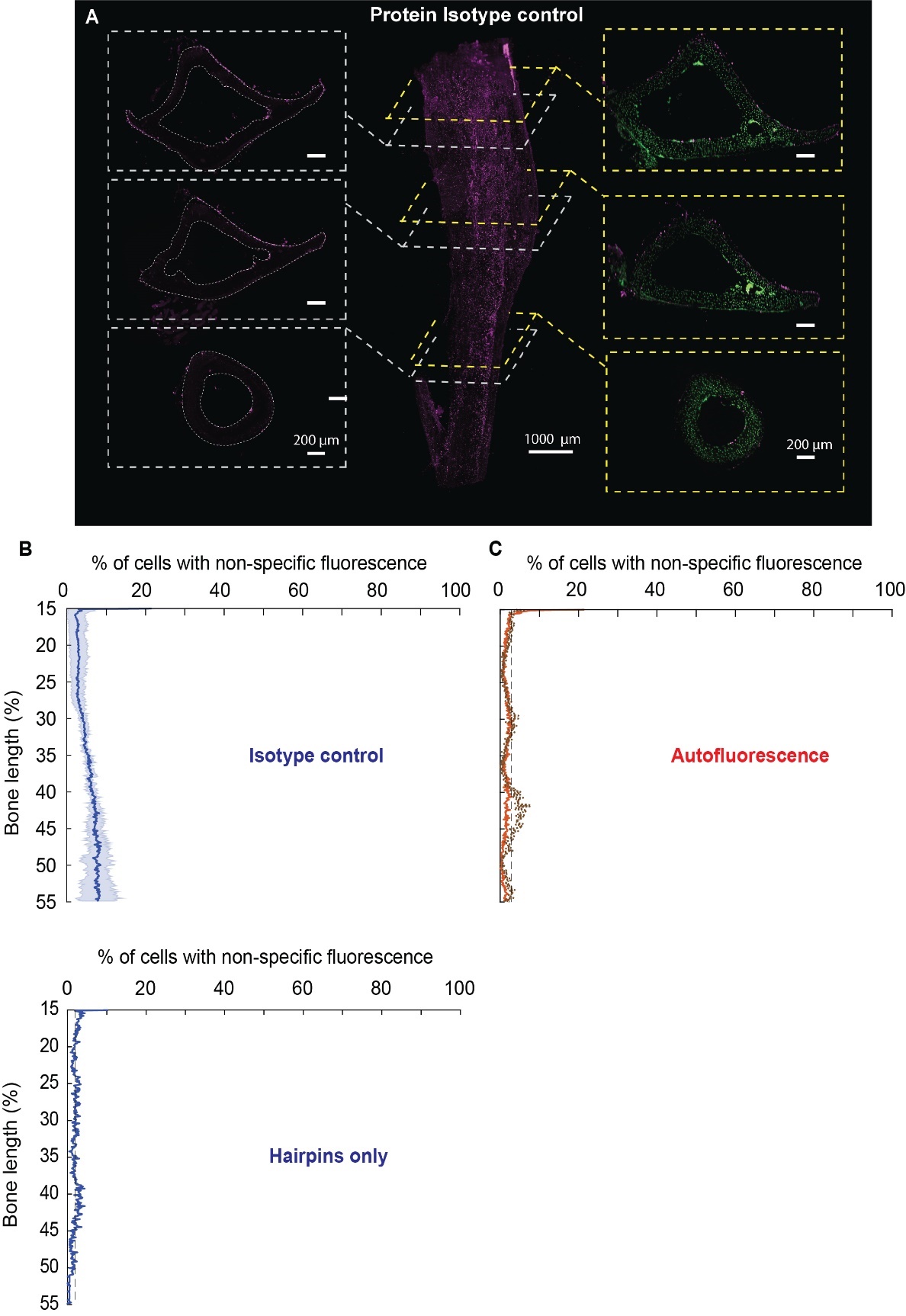

**Supplemental Figure 2:** Spatial investigation of protein negative control samples A) Lightsheet images of a mouse tibia midshaft labeled with Goat IgG Isotype control and secondary antibodies. 3D reconstruction shows some non-specific signal from the midshaft. Virtual cross-sections showed that the non-specific signal only came from the surface of the sample. B) Percentage of cells along the bone length presenting non-specific fluorescence due to incubation with antibodies isotype controls and secondary antibodies (n=3).

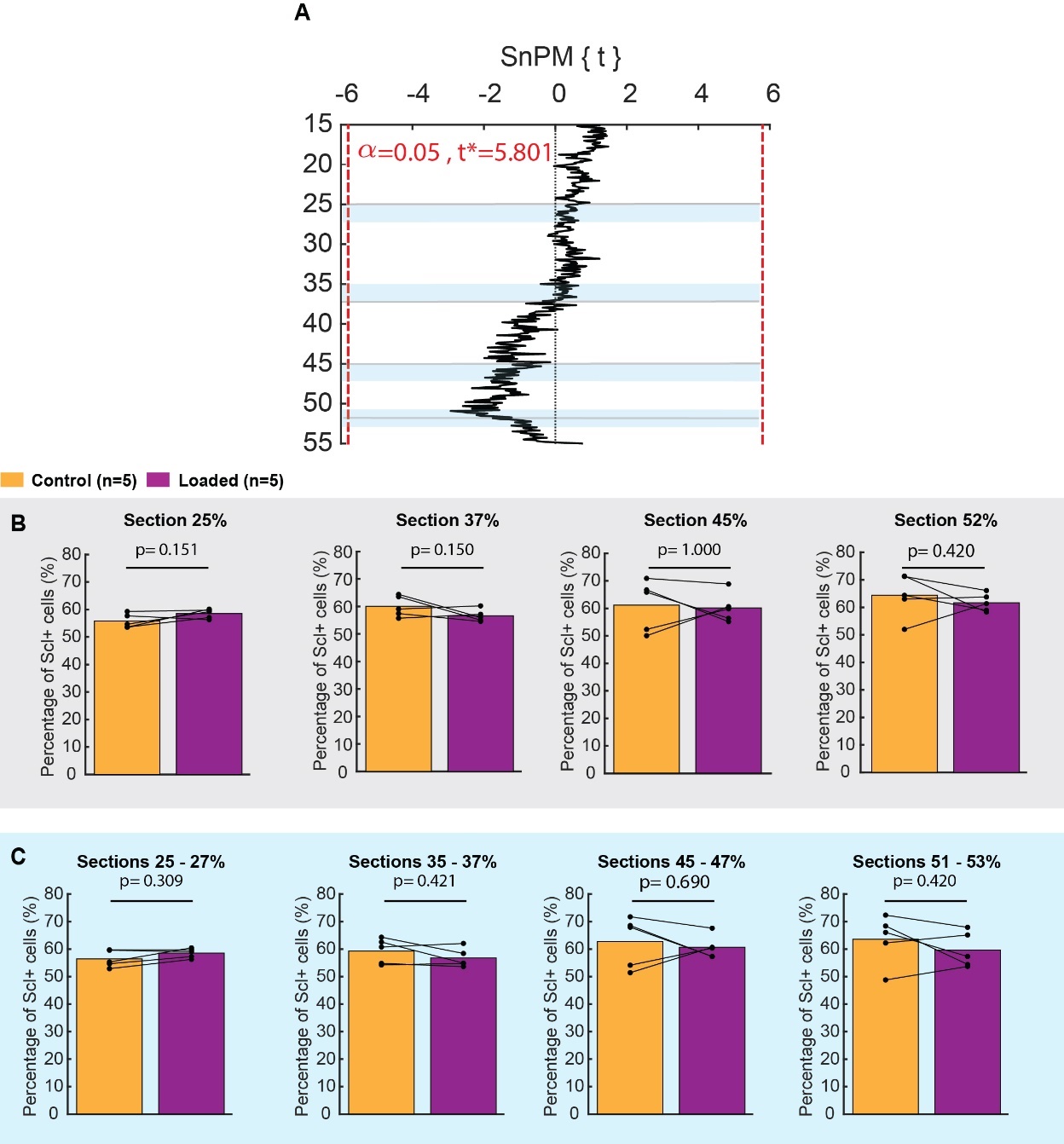

**Supplemental Figure 3:** Non-parametric spm1d two-tailed ttest (α =0.05) testing the significant differences, in percentage of sclerostin-positive cells, between loaded and contralateral legs along the bone length. The black SnPM{t} line corresponds to the statistical test outcomes along the tibia midshaft. A negative SnPM{t} means that the number of positive cells along the loaded legs was lower compared to the contralateral legs. Red dotted lines indicate the critical thresholds (noted t*). When the SnPM{t} curve crosses these critical thresholds, significant differences can be concluded. No significant differences were measured here.

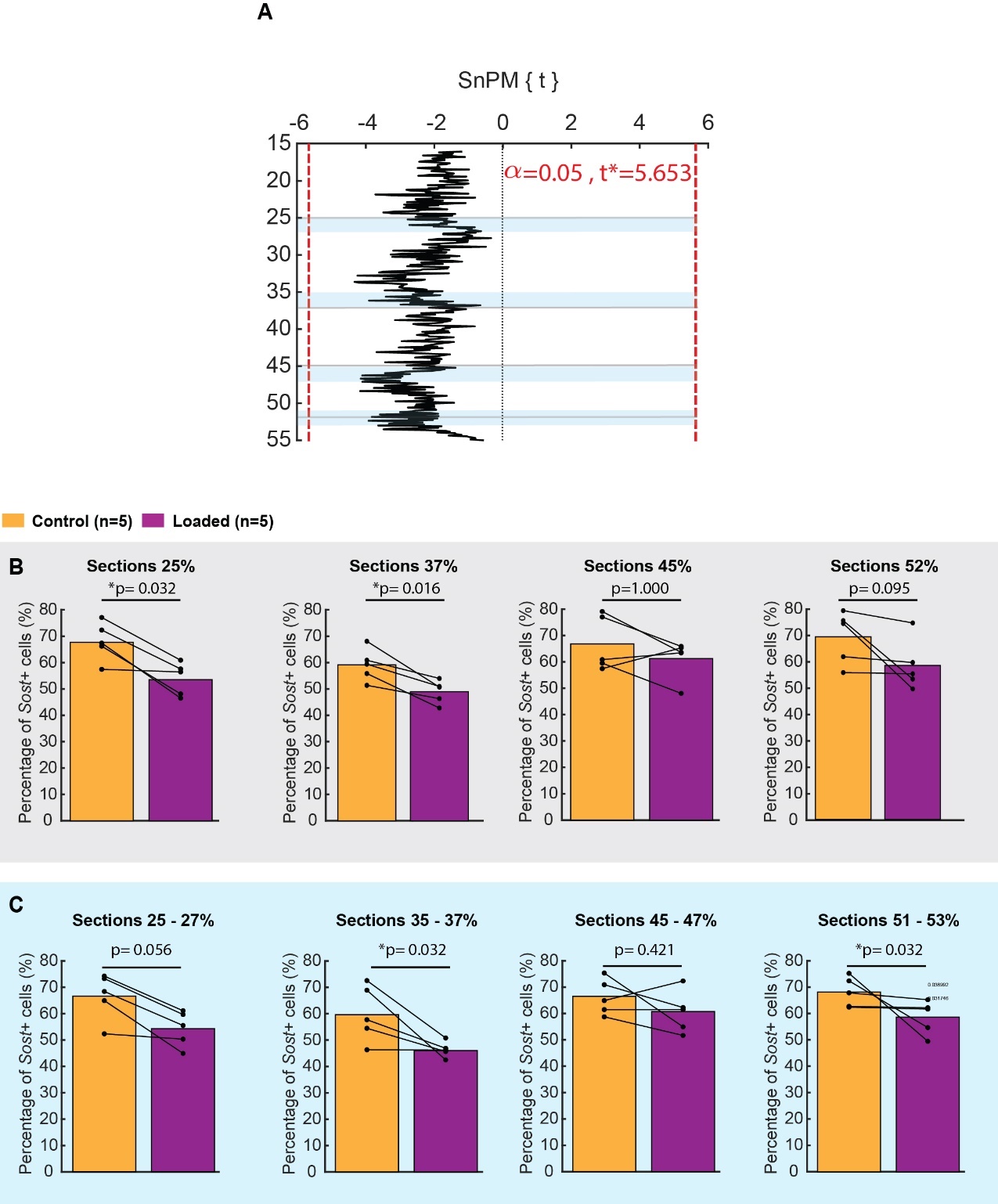

**Supplemental Figure 4:** Non-parametric spm1d two-tailed ttest (α =0.05) testing the significant differences, in percentage of *Sost*-positive cells, between loaded and contralateral legs along the bone length. The black SnPM{t} line corresponds to the statistical test outcomes along the tibia midshaft. A negative SnPM{t} means that the number of positive cells along the loaded legs was lower compared to the contralateral legs. Red dotted lines indicate the critical thresholds (noted t*). When the SnPM{t} curve crosses these critical thresholds, significant differences can be concluded. No significant differences were measured here.
